# Local Fields of the Hippocampus: More than Meets the Eye

**DOI:** 10.1101/008557

**Authors:** Gautam Agarwal

**Affiliations:** Redwood Center for Theoretical Neuroscience, UC Berkeley

## Abstract

A standard methodology in systems neuroscience is to establish correlations between the brain and the world. These correlations have been robustly observed when probing the activity of single neurons. For example, there are cells that activate vigorously when subjects are presented with images of certain celebrities (e.g. the “Jennifer Aniston” neuron, Quiroga et al., 2005). While single neuron measurements provide remarkable insights into the functional specificity of different brain areas, it remains unclear how single cell activity arises from, and contributes to, the collective activity of brain networks. New technologies allow for the simultaneous recording of spikes from hundreds of cells, yet we are still far from observing entire neuronal circuits in action.

An alternative approach in monitoring neuronal populations is to measure local field potentials (LFPs). These potentials arise from the coordinated activity of thousands of nearby neurons. LFPs often exhibit oscillations of various frequencies, which correlate with different states of awareness, such as alertness, relaxation, or drowsiness. However, establishing correlations between LFPs and specific behavioral events can be challenging, because the contributions of different neurons to the LFP usually cannot be distinguished. In this chapter, we examine LFPs recorded from the rat hippocampus. At first glance, these LFPs appear to bear little relation to the subject’s changing environment. However, through appropriate signal processing, we find that carried in the structure of the LFP is a remarkably precise stream of information (Agarwal et al., 2014).

## Activity in the hippocampus

The hippocampus is best known as a region involved in the formation of memory. However, in rodents, it has been most extensively characterized in the context of navigation. As a rat moves through its environment, neurons known as ‘place cells’ activate at specific locations, outside of which they remain largely silent. Different place cells activate at different locations, such that the pattern of activity in a population of cells can be used to estimate the animal’s position (Wilson & McNaughton, 1993). In contrast, the LFP of a navigating rat exhibits a strong, relatively consistent ∼8 Hz rhythm known as theta (Buzsaki & Moser, 2013). It is believed that the theta rhythm arises from the pooled activity of thousands of hippocampal place cells (Geisler et al., 2010); however, because these cells exhibit no known anatomical organization (Redish et al. 2001), the rhythm appears to lack information about the rat’s location. Nonetheless, close observation reveals subtle variations in the relative timing and amplitude of the theta wave, across the spatial extent of the hippocampus (Agarwal et al., 2014). We examined the potential for these micro-variations to carry the information encoded by hippocampal cells about the rat’s location.

## Extracting position information from theta waves

Any signal that is informative about the rat’s position must have two properties. First, every time the rat reaches a position, the message must be the same (reproducibility). Second, different positions must invoke different messages (discriminability) (Borst & Theurnissen, 1999). At first glance, the theta rhythm appears to violate both criteria.

### Reproducibility

Although modulated by the rat’s movement, the theta rhythm largely reflects the internal dynamics of the hippocampal network. Consequently, as the rat reaches a given position, theta will be in an arbitrary phase of its cycle. How does one remove this intrinsically generated source of variability, while preserving the spatio-temporal variations that characterize each theta cycle (as mentioned above)? Here, we take inspiration from radio communication, in which a relatively low-frequency stream of information is embedded in a higher frequency carrier wave. Recovering the signal of interest requires “demodulating” the carrier wave (Fig. 1). When performing a demodulation operation on theta rhythms recorded throughout the hippocampus, one is left with anatomically distributed patterns of relative timing (phase) and intensity (amplitude) that change gradually as the rat moves through space.

**Fig. 1:**
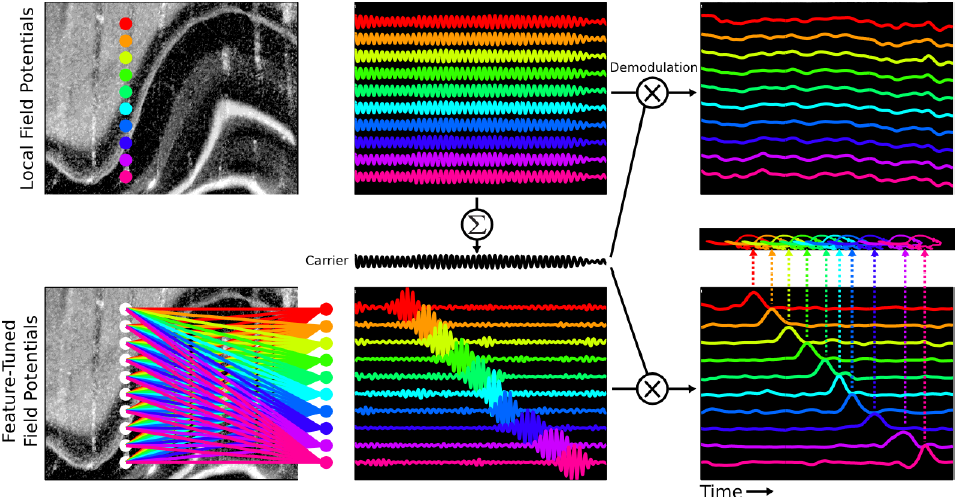
**Top:** The local field potential (LFP) is simultaneously measured from several electrodes in the hippocampus (left), revealing ongoing oscillatory activity (middle). Demodulating the carrier wave common to all electrodes reveals slow modulations of the oscillation (right). **Bottom:** Compressed sensing algorithms can be used to decompose the LFP into distributed patterns (left) that activate sparsely (middle). Removing the carrier wave from the LFP components yields features whose occurrences uniquely predict the animal’s current position (right).

### Discriminability

Another challenge is the apparent lack of selectivity of the theta rhythm. Because the LFP measured at any location sums the electric fields of thousands of neurons with unique place-tuning, it appears to lack place selectivity (Fig 1, top). How can a handful of non-selective readouts be used to reconstruct the information carried by thousands of place-selective cells? Here, we take inspiration from the recently developed theory of compressed sensing (Donoho, 2006), which states that when a signal arises from sparse causes, it can be reconstructed from a subsampled readout (i.e. tens of LFP measurements reflecting the activity of thousands of cells). In our case, since the rat can only occupy a single position at any time, neural activity is evoked by a highly sparse cause. What is more, one can employ learning algorithms to identify LFP components associated with the sparse causes (Bell & Sejnowski, 1995; Olshausen & Field, 1996; Isely et al., 2010). While LFP’s measured at single locations exhibit modest position-dependence, the anatomically distributed LFP patterns identified using these learning algorithms can exhibit exquisite selectivity to the animal’s position (Fig. 1).

Combining the approaches of demodulation and sparse feature learning allows us to conclude that in fact, the theta rhythm changes in a reproducible and discriminable way as the rat moves through its world.

## Mechanisms

We show that the LFP carries precise information about position but make no claim that hippocampal neurons access the LFP signal directly. Although weak electric fields are known to influence the spiking of neurons (“ephaptic coupling” Anastassiou et al., 2011; Reato et al., 2010), this effect may be modest within the behaving brain. The implication of our finding for understanding neural mechanism rests on the fact that the LFP largely reflects the cumulative synaptic input to neuronal population (Buzsaki et al., 2012). Thus, our ability to decode spatio-temporal LFP patterns may hint at how place-tuning arises within a hippocampal neuron as it integrates some 30,000+ synaptic inputs (Megias et al., 2001). More generally, the established practice of recording both LFPs and action potentials within a region can provide a window into how the circuit’s synaptic inputs relate to its spike outputs.

## In Closing

This case study shows how a signal may appear simple, but nonetheless convey a nuanced and rich stream of information. It follows that we should not claim to understand the significance of neural activity unless we understand the language(s) of neurons. A long held view in neuroscience is that neurons can be thought of as feature detectors: high activity in a single neuron signifies the presence of a corresponding feature in the world. An alternative possibility is that a signal’s meaning is context-dependent. For decoding the LFP, we have considered two such contextual factors: the ongoing, internal dynamics of the circuit, and the spatially distributed nature of information. This parallels findings that show at the level of spiking neurons the importance of timing (Laurent, 1996; Brenner et al., 2000) and distributed codes (Schneidman et al., 2011; Riehle et al., 1997).

How does the place code we describe relate to the mnemonic function of the hippocampus? In an accompanying chapter, Walter Freeman suggests that when a percept is stored in memory, it is assigned a corresponding place label in the hippocampus. An implication of Dr. Freeman’s proposal is that associated with the subject’s current position, hippocampal activity should contain traces of its perceptual experience. Yet, we have failed to detect such a form of information. This may be for several reasons. First, the experiments that we analyze are carried out in relatively impoverished settings: the rat runs back and forth on a dark and uniform linear track. It is possible that putting the rat in more ethologically informed environments would enrich the contents of hippocampal activity. Second, our inability to detect perceptual traces might reflect a limitation in analysis. We considered exclusively the theta rhythm, although the hippocampus contains many other rhythms, such as gamma (25-140 Hz, Colgin & Moser, 2010). It is possible that while patterns in theta can be decomposed using algorithms that exploit sparsity, other forms of learning will be more appropriate in dissecting the structure of other oscillations in the LFP. Ultimately, a major current limitation in our approach is that the patterns we identify must be validated by correlating them to externally measurable observables (in our case, the rat’s position). More sophisticated methods are needed to relate neural activity to the pristine contents of inner experience (Hurlburt, 2013).

## Acknowledgements

I would like to thank my advisor Friedrich Sommer and collaborator-advisor Gyorgy Buzsaki for their feedback and insights throughout the course of this project, as well as members of the Redwood Center and Buzsaki lab for discussions. This work was supported by NIH National Research Service Award #1F32MH093048. The data used in this project is available at crcns.org.

